# Nanopore Fingerprinting of Supramolecular DNA Nanostructures

**DOI:** 10.1101/2022.06.23.496947

**Authors:** Samuel Confederat, Ilaria Sandei, Gayathri Mohanan, Christoph Wälti, Paolo Actis

## Abstract

DNA nanotechnology has paved the way for new generations of programmable nanomaterials. Utilising the DNA origami technique, various DNA constructs can be designed, ranging from single tiles to the self-assembly of large-scale complex multi-tile arrays. These DNA nanostructures have enabled new applications in biosensing, drug delivery and other multifunctional materials. In this study, we demonstrate real-time, non-destructive and label-free fingerprinting of higher-order assemblies of DNA origami nanostructures using solid-state nanopores. Using this approach, we quantify the assembly yields for each DNA origami nanostructure with single-entity resolution using the nanostructure-induced charge introduced in the nanopore as a discriminant. We compare the assembly yield of the supramolecular DNA nanostructures obtained with the nanopore with agarose gel electrophoresis and AFM imaging and demonstrate that the nanopore system can provide enhanced information about the nanostructures. We envision that this nanopore detection platform can be applied to a range of nanomaterial designs and enable the analysis and manipulation of large DNA assemblies in real-time with single-molecule resolution.

**STATEMENT OF SIGNIFICANCE:** We demonstrate a single molecule high-throughput approach for the analysis of higher-order DNA origami assemblies with a crowded nanopore. The technique enables the characterisation of DNA origami nanostructures at statistically relevant numbers in real-time and at single-molecule resolution while being non-destructive and label-free, and without the requirement of lengthy sample preparations or use of expensive reagents. We exemplify the technique by demonstrating the quantification of the assembly yield of DNA origami nanostructures based on their equivalent charge surplus computed from the ion current signals recorded. Compared to the standard analysis methods of AFM and agarose gel electrophoresis, the nanopore measurements provides enhanced information about the nanostructures.

## INTRODUCTION

The use of DNA as a building block for the engineering of nanoscale materials is one of the corner stones of DNA nanotechnology (1). In particular, the invention of the DNA origami approach, which exploits Watson-Crick base-pairing between a single-stranded DNA scaffold and multiple short staple-strands to fold the scaffold into a specific predefined geometry (2), allowed the folding of nanostructures with large surface area while simultaneously enabling spatially controlled assemblies and site-specific chemical functionalisation (3). These unique characteristics have enabled the assembly of nanostructures and patterns with controlled geometry and function, either by directly folding the scaffold (4, 5), or via higher-order assembly of pre-formed DNA tiles (6-8), which found applications in biosensing (9-13), drug delivery (14, 15), and as tools for studying biological processes (16, 17), *inter alia*. Fuelled by these rapid developments in engineering and fabricating complex DNA nanostructures, there is an increasing demand for their characterisation, including the assessment of assembly yields. Traditionally, the characterisation of DNA constructs has relied on methods such atomic force microscopy (AFM), agarose gel electrophoresis, and transmission electron microscopy (TEM) (18). In recent years, nanopores have been established as a powerful tool for the characterisation of DNA nanostructures with single molecule resolution. Nanopore sensing is based on the measurement of transient perturbations of the ion current through a nanopore caused by the translocation of an analyte (19, 20). The characteristics of these perturbations, such as the amplitude and duration, provide information about the physico-chemical properties of the translocating molecule, including size, charge, and shape (21, 22). Nanopore sensing has also been successfully applied for the detection of colloidal nanoparticles (23, 24) and biological nanoparticles including virus particles (25), ribonucleic particles (12), protein aggregates(26), and DNA nanostructures (27-30). Our group has previously demonstrated the analysis of 2D DNA origami (31) and single molecule biosensing (13) using nanopores and has recently shown the marked enhancement of single molecule detection within a crowded nanopore (32). Other groups used DNA-based nanoswitches for sensing applications using nanopores (33), and a similar approach has been used for miniaturised molecular data storage (34).

Here, we report the single molecule detection of supramolecular DNA assemblies by solid-state nanopore analysis. We demonstrate that the perturbations in the ion current caused by the translocation of a DNA origami nanostructure can fingerprint different states of higher-order assemblies, ranging from an individual monomer building block to multimer assemblies. We quantify the assembly yields of a range of higher-order assemblies of DNA nanostructures with single-entity resolution and benchmark the nanopore analysis against agarose gel electrophoresis and AFM imaging.

## MATERIALS AND METHODS

### DNA nanostructures

The design of the four DNA nanostructures used here follows the design published by Tikhomirov *et al*. (*6, 35*) and was carried out using the CaDNAno 2.3 software (36). All four nanostructures were folded from the single-stranded (ss)DNA M13mp18 (7249 bp) following the standard procedure for DNA origami fabrication (4). For the higher-order assemblies, the individual tiles were first folded individually using three main kinds of staples: bridge staples, interior staples, which constitute the main body of the tile, and a specific set of edge staples, which allow for a specific interaction between the structures (the complete list of the staples is provided in the SI file). While the same set of bridge staples and interior staples is shared by all DNA nanostructures, each one of them has a different set of edge staples (Figure S14). The edge staples can be “giving” (featuring a two-nucleotide extension), “receiving” (with a two-nucleotide truncation) or inert (characterised by two hairpins, forming a loop conformation, and preventing the further higher-order assembly of DNA origami along that side). Each structure can have eight giving/receiving staples or five inert staples. The use of five inert staples instead of eight as in the case of giving and receiving staples was justified by Tikhomirov *et al*. to prevent the deformation of the tiles and undesired interactions, seen when using a full set of eleven hairpins (35). For the assembly, we followed published protocols (6, 35) where the edges of each tile can be labelled as north, south, east, and west. Each set of interactive edge staples is complementary to just one other specific set of edge staples (north giving edges are complementary to west receiving edges, west giving edges to south receiving edges, east giving edges to north receiving edges, and south giving edges to east receiving ones), allowing for a targeted assembly and avoiding spurious interactions. To fold the structures, the individual monomer structures having complementary edge sequences were mixed together in equal concentrations and volumes, in order to increase the yield of the assembly. Specifications on the sequences and the interactions between the pre-assembled monomers for all the higher-order structures used in this work are provided in the SI file (36).

### DNA nanostructures folding

The single-stranded M13mp18 DNA scaffold 7249 bp was purchased from New England Biolabs (NEB, UK) at an initial concentration of 100 μM in 1xTE buffer, pH 8 (Sigma Aldrich, UK). The staple strands were purchased from Integrated DNA Technologies (IDT, UK) and resuspended in 1x TE buffer pH 8 to a final concentration of 100 μM. The negation strands were purchased from IDT and resuspended in 1x TE buffer pH 8 to a final concentration of 200 μM. For the folding, the ssDNA scaffold (final concentration 10 nM) was mixed with the staple strands (final concentration 75 nM) in 1x TE buffer pH 8 with 12.5 mM of MgAc (Sigma Aldrich) in 80 μL total volume. To fold the individual tiles, the solution of scaffold and staples was heated to 90°C for 2 min and annealed using a temperature ramp from 90°C to 20°C at 6 sec per 0.1°C in a Mastercycler Nexus PCR Thermal Cycler (Eppendorf). Following the annealing step, the negation strands were added to the solution to achieve a final concentration of 375 nM in 80 μL final total volume. The addition of the negation strands, which are complementary to the edge staples, inactivates the excess edge staples so the monomers can be used in higher-order assemblies (6, 35).

Another temperature ramp was then applied, going from 50°C to 20°C at 2 sec per 0.1°C. The folded structures were purified using Sephacryl S400 (GE Healthcare, UK) size exclusion columns in order to remove the excess staple strands, and the product was eluted in 1x TE pH 8, 12.5 mM MgAc. The concentration of the individual tiles was measured using a NanoDrop 2000c Spectrophotometer (Thermo Scientific, UK) to prepare solutions with equal concentration. To fold the higher-order assemblies, equal quantities of the required individual monomers were mixed in PCR tubes and annealed using a temperature ramp from 55°C to 45°C (at 2 min per 0.1°C) followed by a ramp from 45°C to 20°C (at 6 sec per 0.1°C). The folded nanostructures were imaged by AFM to confirm the successful assembly.

### Atomic Force Microscopy (AFM) Imaging

For AFM imaging, 10 μL of purified DNA sample diluted to a final concentration of 0.5 nM in 1x TE pH 8, 12.5 mM MgAc were deposited on a freshly cleaved mica substrate (Agar Scientific, UK) and incubated at room temperature for 15 min. Additional 150–180 μL of 1x TE pH 8, 12.5 mM MgAc buffer was added to the sample to facilitate the imaging. The samples were imaged using a Dimension Fastscan Bio (Bruker, UK) in tapping mode in liquid with Fastscan D Si_3_N_4_ cantilevers with a Si tip (Bruker, UK). We used the following imaging parameters: scan rate = 2–8 Hz, 256 samples/line, amplitude setpoint = 150–300 mV, drive amplitude = 3000 mV, integral gain = 1, proportional gain = 5. The data were processed using Nanoscope analysis 1.9.

### Agarose gel electrophoresis

The quality of the DNA origami nanostructures was further inspected with agarose gel electrophoresis (Figure S2). For this, 50 mL 0.7% agarose gel was prepared using 1x TAE buffer containing 12.5 mM Mg(Ac)_2_. The DNA origami samples were prepared by mixing a 20 μl aliquot at a concentration of 10 ng/μl with 4 μl of 6x Tri Track Loading Dye (Thermo Scientific, UK). M13mp18 scaffold was added as a reference. A GeneRuler 1kb DNA double-stranded ladder (Thermo Scientific) was used as molecular marker and positive control (Figure S2). The running buffer consisted of 1x TAE and 12.5 mM Mg(Ac)_2_. The gel was run at a constant voltage of 70 V for 120 min. The gel was then stained for 30 min with Diamond Nucleic Acid Dye (Promega) diluted in 1x TAE buffer. For this, 5 uL 10,000x concentrated Diamond Nucleic Acid Dye was diluted in 1x TAE buffer and the gel incubated at room temperature for 30 min on a rocking platform. The agarose gel imaging was carried out using the GeneSnap software. Quantification of the gel bands for each DNA origami sample was done using ImageJ.

### Nanopore fabrication and characterisation

The nanopores were fabricated starting from 1.2 mm x 0.9 mm quartz capillaries (QF120-90-10; Sutter Instrument, UK) with the SU-P2000 laser puller (World Precision Instruments, UK), using a two-line program: (1) HEAT, 800; FILAMENT, 4, VELOCITY, 30; DELAY, 145’ PULL, 95; (2) HEAT, 730, FILAMENT, 3; VELOCITY, 40; DELAY, 135; PULL, 150. The pulling parameters are instrument specific and lead to nanopipettes with a nanopore diameter of approximately 160 nm. The nanopipettes were characterized by measuring their pore resistance in 0.1 M KCl (∼40 MΩ) and the nanopore dimensions were confirmed by Scanning Electron Microscopy (SEM) using a Nova NanoSEM at an accelerating voltage of 3–5 kV (Figure S1).

### Nanopore translocation measurements

The translocation experiments were carried out by filling the nanopipette with the translocation buffer (100 mM KCl, 0.01% Triton-X, 10 mM Tris, 1 mM EDTA, pH 8.0) containing the DNA origami sample at a concentration of 500 pM. The nanopipette was then immersed in a 100 mM KCl bath with the addition of 50% (w/v) Polyethylene Glycol (PEG) 35 kDa (ultrapure grade, Sigma Aldrich). An Ag/AgCl wire (0.25 mm diameter, GoodFellow UK) was inserted in the nanopipette barrel and acted as the working electrode, while a second Ag/AgCl wire was immersed in the bath and acted as the counter and reference electrodes. The DNA origami nanostructures were driven from inside the nanopipette into the external bath by applying a negative potential to the working electrode placed inside the nanopipette with respect to the reference electrode in the bath. The ion current was recorded with a MultiClamp 700B patch-clamp amplifier (Molecular Devices) in voltage-clamp mode. Data was acquired at a 100 kHz sampling rate with a 20 kHz low-pass filter using the pClamp10 software (Molecular Devices). The ion current traces were further analysed with the MATLAB script Transanlyser, developed by Plesa *et al*. (37). The obtained translocation events were analysed by applying a 7-sigma threshold level from the baseline, and only the events above the threshold were considered as translocation events (Figure S6). The obtained events were further analysed and plotted using Origin 2019b.

## RESULTS AND DISCUSSION

### DNA nanostructures assembly

In this study, we demonstrate the discrimination of supramolecular DNA nanostructures with solid-state nanopores by exploiting the unique signature in the translocation signal resulting from the perturbation when the nanostructures pass through the nanopore (Figure 1). We assembled four different DNA nanostructures, starting from a pre-assembled DNA origami (85 nm x 85 nm) used as a building block (referred to hereafter as monomer), followed by a two-monomer assembly (dimer), a three-monomer assembly in an L-shape (trimer), and lastly a 2×2 array of monomers (referred to hereafter as 2×2) (Figure 2A). AFM measurements confirmed the correct folding of the DNA nanostructures (Figure 2B).

**Figure 1:**
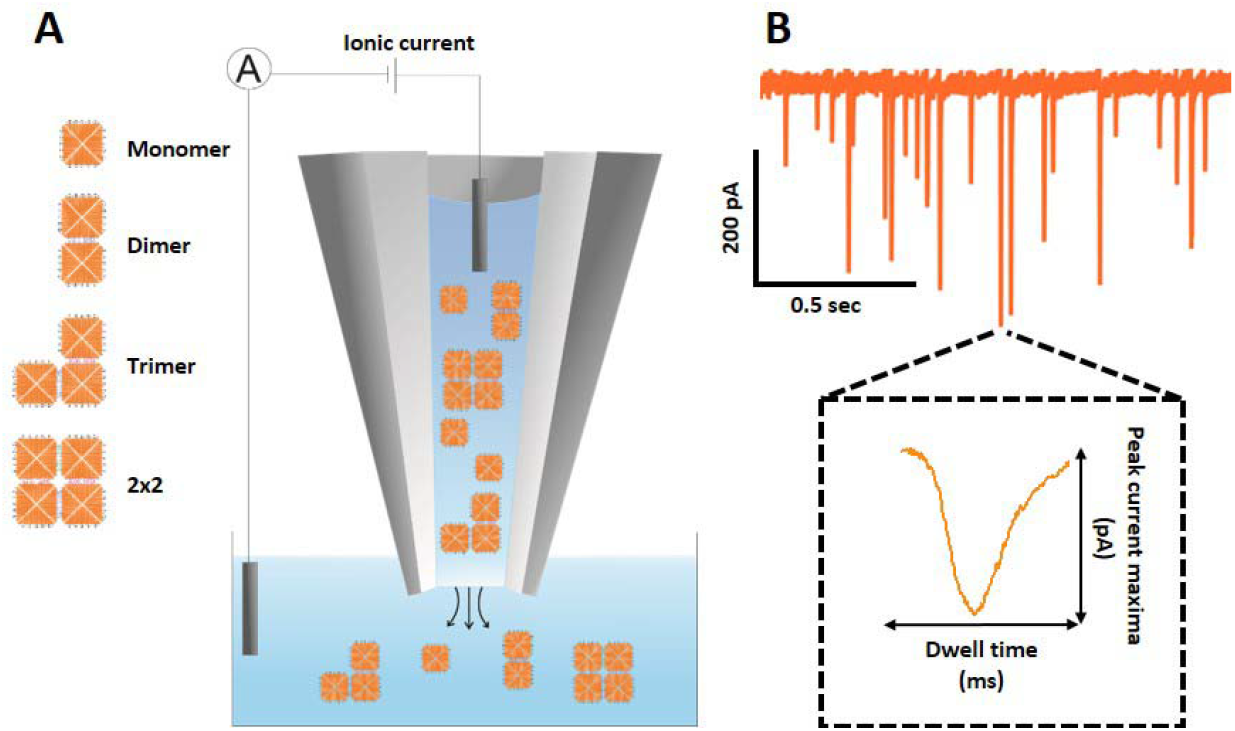
**(A)** Schematic representation of the nanopore setup. The DNA origami nanostructures are translocated from inside the nanopore into the outer bath upon the application of a negative potential bias while the ion current is measured. **(B)** Representative ion current trace showing translocation events. A representative event is show in the inset.

**Figure 2:**
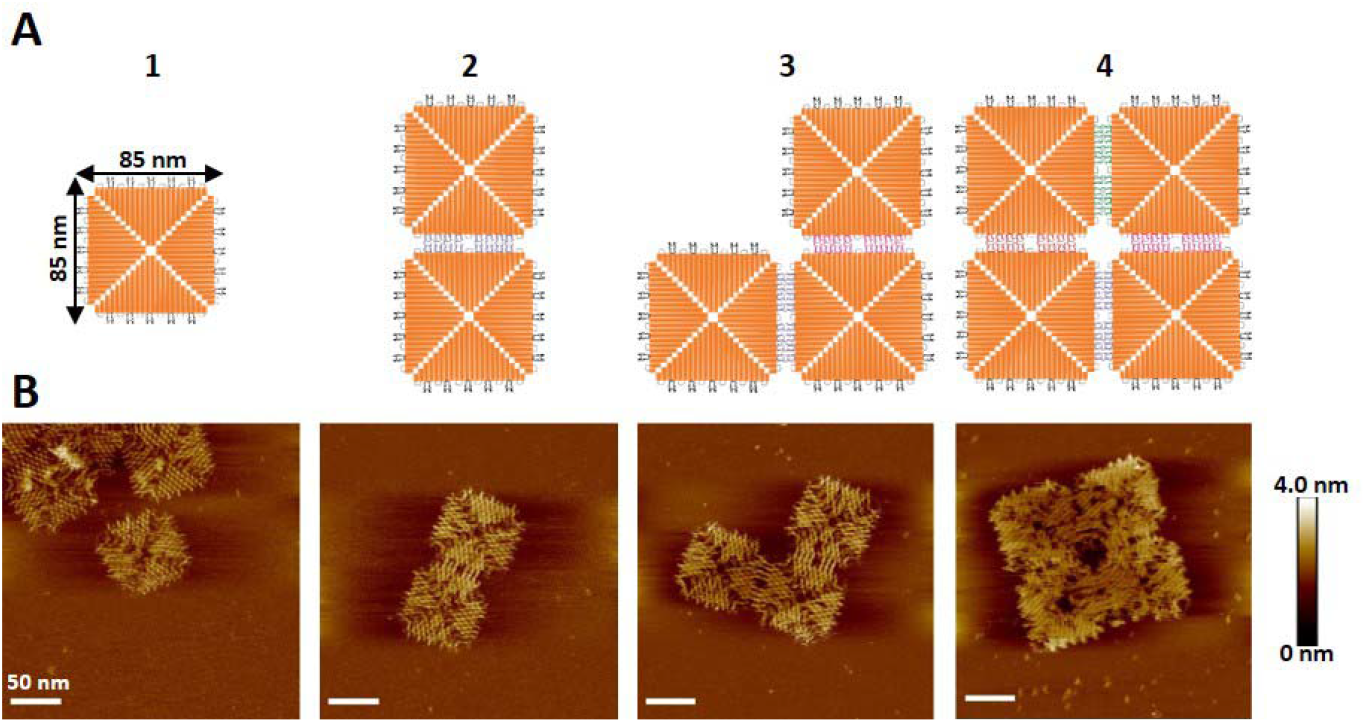
**(A)** Schematic representation of the DNA nanostructures, starting with a single monomer building block (1), dimer (2), trimer (3), and 2×2 (4). **(B)** AFM micrographs of the DNA nanostructure depicted in panel A (50 nm scale bar).

### Nanopore analysis of DNA nanostructures

We chose a nanopore diameter of 160 nm to be large enough to accommodate the largest DNA nanostructure (2×2) while retaining a sufficient signal-to-noise ratio for the detection of the monomer. The detection of our smallest (monomer) and largest (2×2) DNA nanostructures with a fixed pore size was facilitated by adapting the bath conditions where the nanopipette is immersed. We have previously demonstrated the marked enhancement in single-molecule-detection sensitivity when the commonly used macromolecular crowder polyethylene glycol (PEG) is added to the bath solution to a final concentration of 50% w/v (32). The presence of PEG resulted in translocation peaks that were well-resolved from the ion current baseline (Figure 3A).

**Figure 3:**
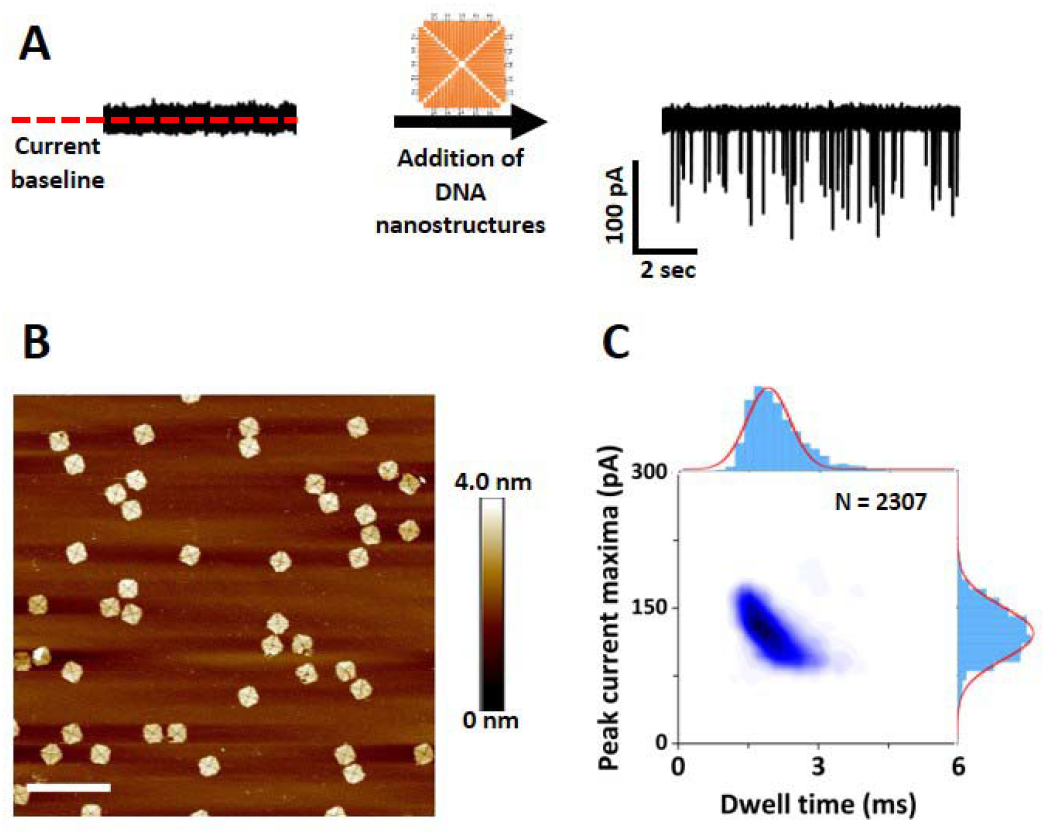
**(A)** Ion current trace before and after the addition of the DNA origami monomer. **(B)** AFM micrograph of a monomer DNA origami sample (400 nm scale bar). **(C)** Density scatter plot of the monomer DNA origami sample with the peak current maxima *versus* dwell time and their corresponding marginal histograms. The solid lines in the marginal histograms represent a Gaussian distribution fitted to the data. A total of *N* = 2307 translocation events were analysed, and their corresponding peak current maxima and dwell time values extracted.

In contrast, the translocation of monomer DNA nanostructures in the absence of PEG resulted in a low signal-noise-ratio, constraining the detection of the monomer nanostructures (Figure S4). Furthermore, the presence of PEG allowed us to detect all DNA nanostructures with a high signal-noise-ratio and to record stable ion current traces with high capture rate (∼5 events/sec at -300mV) for several minutes without any evidence of nanopore clogging (Figure S5). We first investigated the translocation of the monomers (Figure 3B) from inside the nanopipette into the outside bath under a constant voltage bias of -300 mV (Figure 3A). The translocation of the monomer DNA nanostructures resulted in conductive (current enhancing) translocation signal peaks, where each peak is associated with the passage of one molecule. No peaks are detected under the same conditions if no analyte was added to the nanopipette (Figure S7A,B). The translocation events of the monomer sample can be characterised using the peak current maxima (maximum amplitude of the peak from the baseline) and dwell time (duration time of an event). The results (*N* = 2307 peaks) are displayed in the density scatter plot of Figure 3C, which shows that the translocation peaks fall within a well-defined area. The histograms showing the peak current maxima and the dwell time distributions, respectively, are also shown in the figure. Both distributions have been fitted with a Gaussian distribution function yielding an average peak amplitude of 123±27 pA and an average dwell time of 2.1±0.6 ms. Upon increasing the applied voltage, we observed an increase in the peak current maxima of the conductive peaks, as well as a decrease of the dwell time (Figure S8A-D), suggesting that the DNA nanostructures are electrophoretically driven through the pore. This observation was also confirmed by translocation controls where no potential was applied (Figure S7D) or the applied bias was reversed (Figure S7C). Furthermore, we also probed the effect of DNA nanostructure concentrations and the effect of applied voltage on the translocation event frequency. With an increasing concentration of DNA nanostructures, more molecules are expected in the capture region of the nanopore which results in a larger number of translocation events. We confirmed this observation by running translocation measurements at different concentrations (50-500 pM), as shown in Figure S8F. Moreover, increasing the applied voltage induces a larger capture region, leading to a greater number of translocation events (Figure S8E). These observations are consistent with established physical models reported in the literature on the study of nucleic acids translocations through solid-state nanopores (38-40). We then analysed the higher-order-assembly DNA origami samples, using the same translocation conditions as for the monomer analysis.

Figure 4A depicts the current versus time traces obtained for the monomer and 2×2 samples where a new population of translocation events can be observed for the 2×2 sample. While the monomer sample led to translocation events with peak current maxima of less than ∼150 pA, events with significantly larger current amplitudes can be seen for the 2×2 sample – the current range within which these new peaks fall is indicated by the blue shading. This observation suggested that our nanopore platform is able to discern the two ‘extreme’ designs of the higher-order DNA origami assemblies introduced in Fig 2A using a fixed nanopore diameter. The density scatter plots in Figure 4B show a difference in the clustering of the translocation events between the monomer and the 2×2 sample. One of the clusters for the 2×2 sample is very similar to the single population observed in the monomer sample. However, the 2×2 sample shows an additional broader and less well-defined cluster centred at around 4 ms and 250 pA, which is also evidenced by the distributions plots in Figure 4C. The distributions were fitted with a single and a two-peak Gaussian distribution (shown as solid lines in the plot). While the monomer sample yielded an average peak current maximum of ∼123 pA (see also Figure 3C), the distribution corresponding to the more well-defined cluster in the 2×2 assembly sample density scatter plot which is reminiscent of the one found for the monomer yielded an average peak current maximum of ∼170 pA, and the broader less well-defined distribution an average peak current maximum of ∼260 pA. Despite the slight difference in average peak currents, we hypothesised that the more well-defined cluster of the 2×2 sample corresponds to monomers in the sample which were not successfully assembled into higher-order assemblies. We confirmed this hypothesis by spiking the 2×2 sample with increasing concentrations of the monomer. As can be seen in the Figure S10, we observed a shift towards the monomer cluster when increasing the number of monomers spiked into the 2×2 sample. However, it is unlikely, given the relatively broad width of the distribution, that the less well-defined cluster represents only a single assembly state. This is further supported by both AFM imaging (Figure S3) and gel electrophoresis (Figure 6D) which suggest that the 2×2 sample did not only contain the 2×2 and monomer nanostructures, but also other assembly intermediates. In order to deconvolute further the broad less-well defined cluster of translocation events in the 2×2 sample we analysed all four DNA origami assembly samples, and the respective density scatter plots are shown in Figure 5.

**Figure 4:**
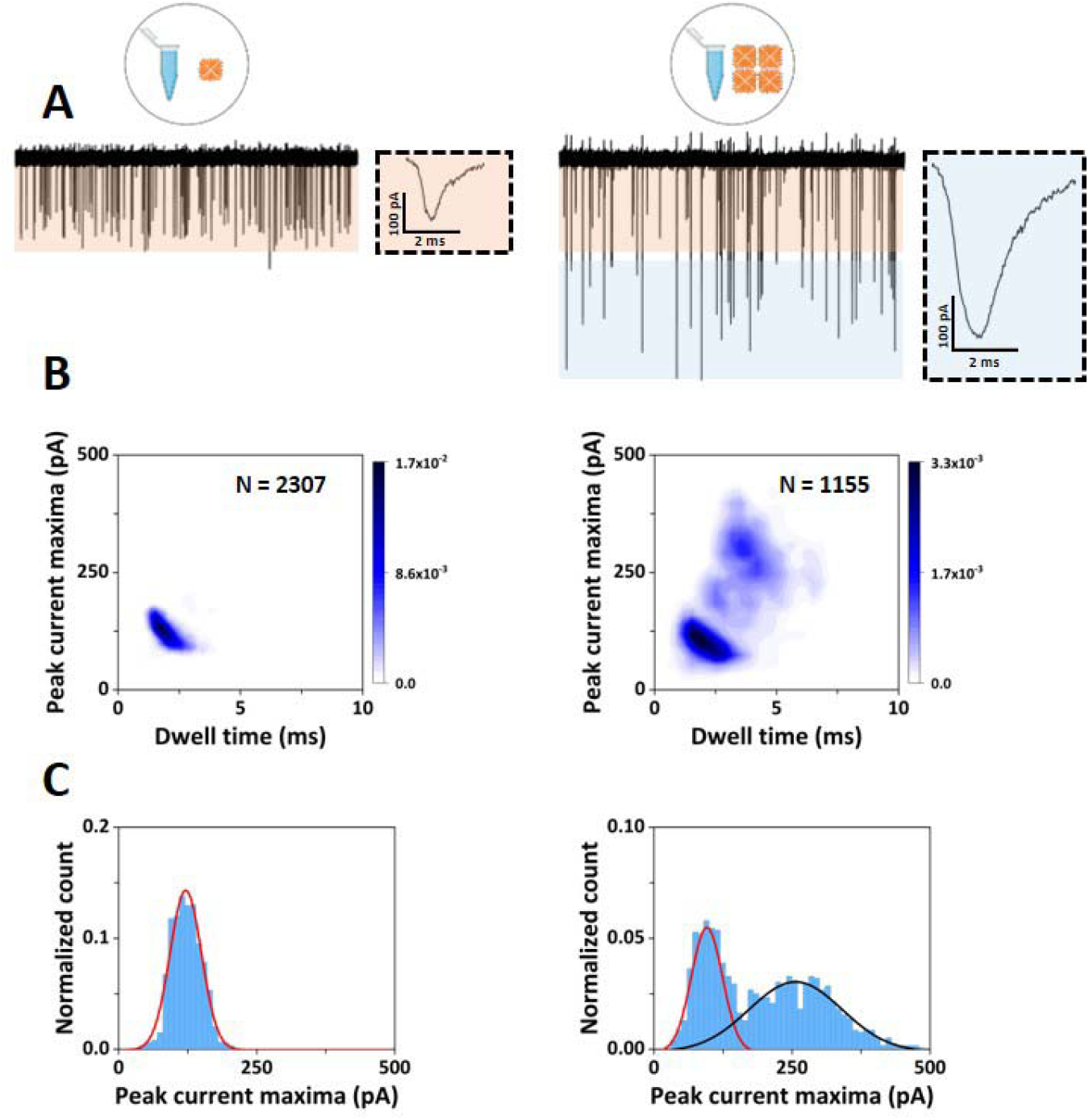
**(A)** Nanopore translocation current traces of the monomer (left) and the 2×2 (right) samples. The current and time scales for the two traces are identical. The orange shading indicates the current range <150 pA, and the blue shading indicates the current range >150 pA. The insets show a representative event for each sample. **(B)** Density scatter plots of peak current maxima as a function of dwell time for the monomer sample (left) and the 2×2 sample (right). **(C)** Histograms of the current peak maxima for the monomer sample (left) and the 2×2 sample (right). The solid lines represent Gaussian fits to the data.

**Figure 5:**
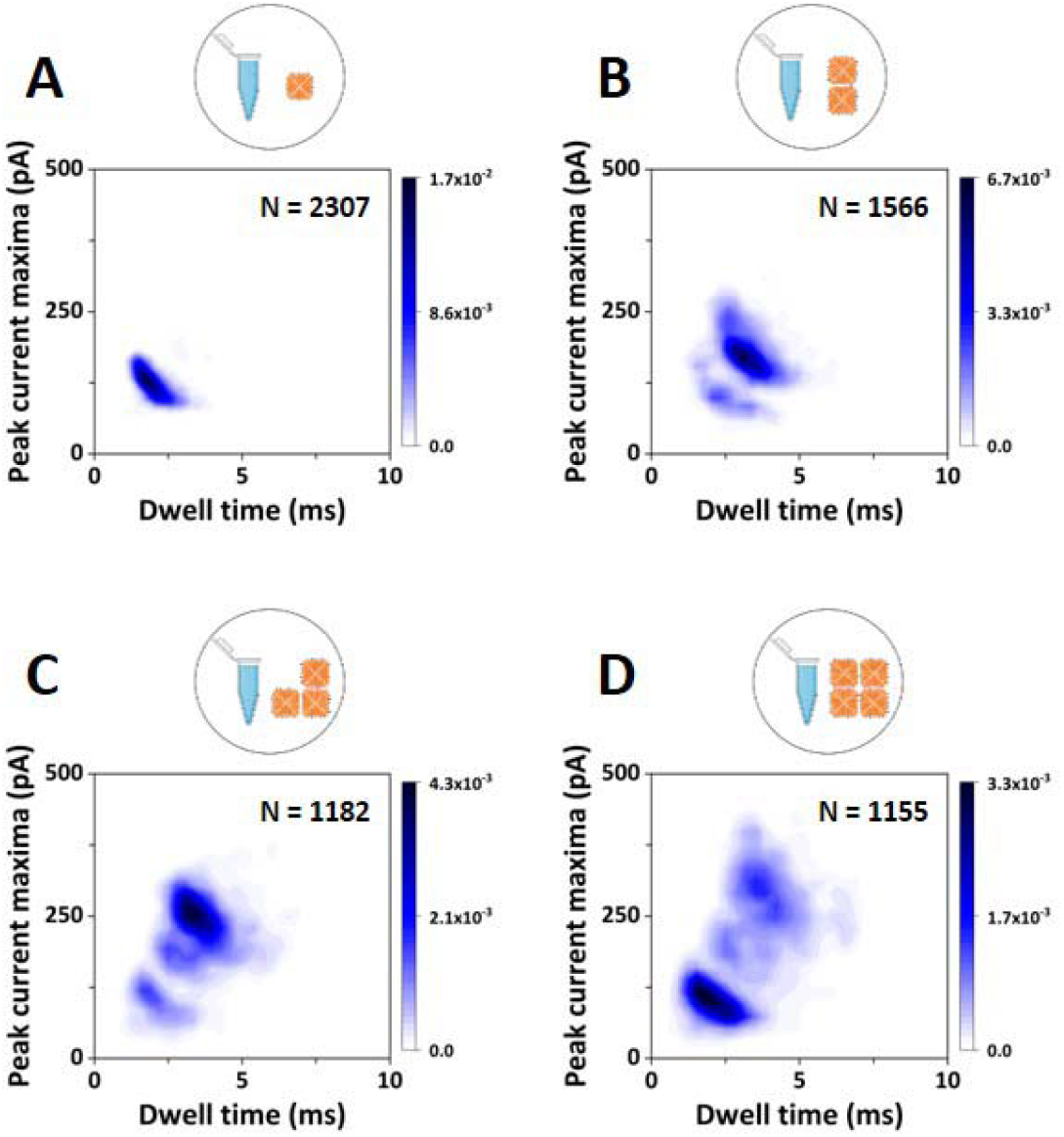
Density scatter plots of peak current maxima as a function of dwell time for the monomer sample **(A)**, dimer sample **(B)**, trimer sample **(C)** and 2×2 sample **(D)**. The number of events analysed, *N*, for each sample is given in each panel.

**Figure 6:**
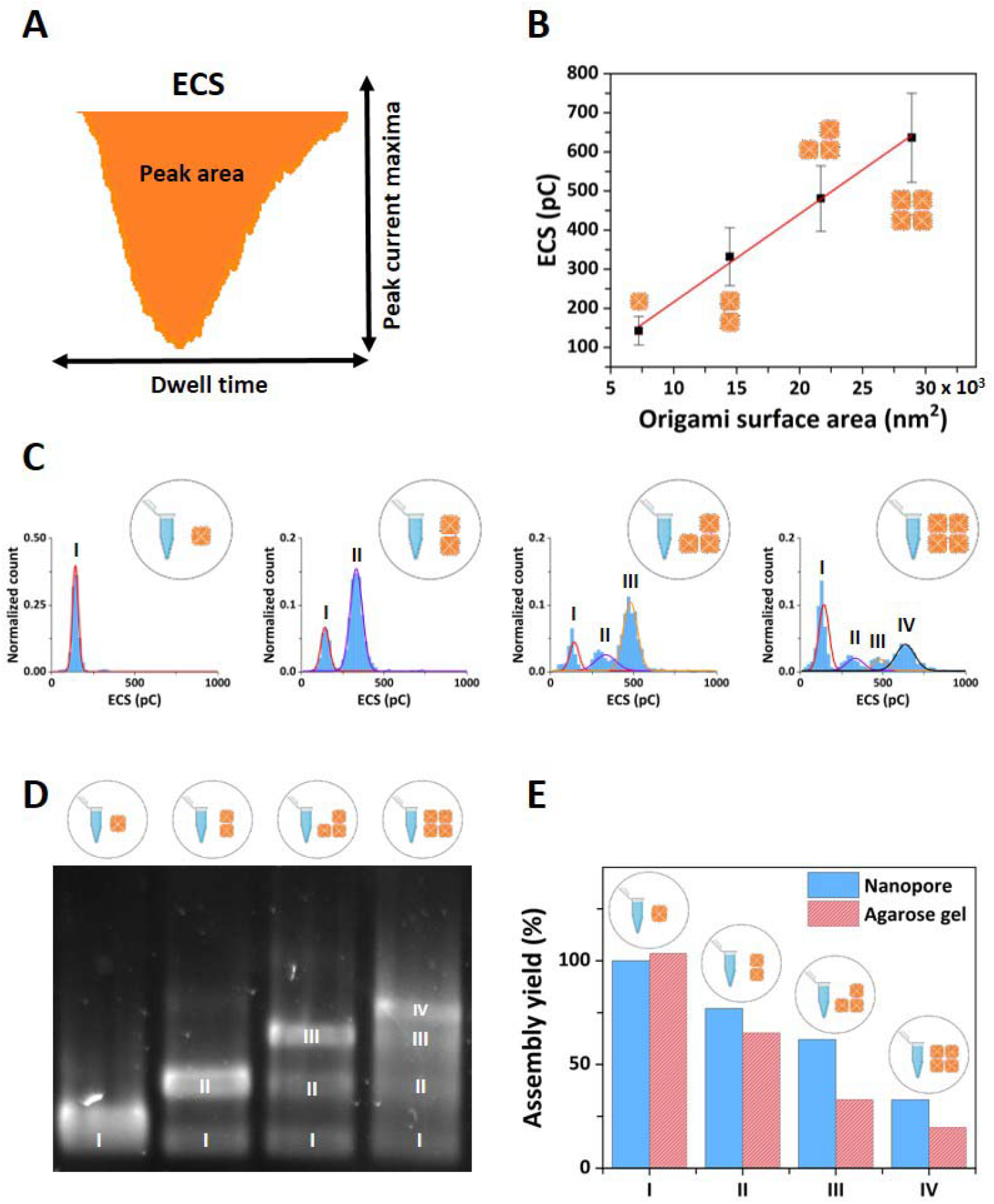
**(A)** Schematic representation of the concept of Equivalent Charge Surplus (ECS) which is obtained from calculating the area of the translocation peaks. **(B)** ECS as a function of the DNA origami surface area for the four DNA nanostructure assembled (the red line represents a linear fit to the data). The error bars represent the width of the Gaussian fits displayed in panel C. **(C)** ECS histograms of the DNA origami samples; from left to right: monomer sample, dimer sample, trimer sample, and 2×2 sample. The distributions were fitted with single or multi-peak Gaussian distributions. The notation I-IV marks the peaks corresponding to each of the DNA nanostructures in the respective sample (*e*.*g*. the dimer sample shows two peaks in the ECS distribution, attributed to the presence of the monomer (I) and dimer DNA origami (II) in the sample). **(D)** Agarose electrophoresis gel of the DNA origami samples; from left to right: monomer samples, dimer sample, trimer sample, and 2×2 sample. The I-IV notation of the gel bands marks the presence of the nanostructure component in the respective sample, similar to the notation used for the ECS distribution in panel C of the figure. **(E)** Bar chart comparison of the assembly yield of each DNA nanostructure (blue bar: nanopore data, red bar: gel electrophoresis data).

All four DNA origami assembly samples (monomer, dimer, trimer, 2×2) were analysed individually with the nanopore platform using the same translocation conditions. The density scatter plots allowed us to identify different clusters for each sample, and for all higher-order assembly samples (Figures 5B-5D) we noticed a consistent presence of the monomer cluster. This is not unexpected – the higher order assembly processes have finite yields and a certain percentage of unassembled monomers are expected to remain. Furthermore, the higher-order assembly is reversible, and while the assembled construct is expected to be energetically favourable, dissociation of higher-order assemblies occurs over time. The translocation peak characteristics of each DNA origami sample showed differences in terms of their peak current maxima and dwell time distributions (Figure S9). The dimer sample yielded one additional well-defined cluster of peaks, with slightly higher peak current maxima and dwell time averages, consistent with what is expected for the translocation of assembled dimers. However, the situation is much less clear for the trimer sample – similar to the 2×2 sample – where in addition to the well-defined cluster originating from the monomer peak an additional number of much less-well defined clusters are observed. While the clustering is clearly different between the four samples, it would be highly desirable to be able to distinguish the different assembly intermediates within a sample.

### DNA origami assembly yield analysis

Finally, in addition to distinguish different assembly intermediates within a mixed sample, we also aimed to go beyond the standard analysis of DNA origami nanostructures assembly (AFM and gel electrophoresis) and quantify the percentage of each DNA nanostructures present in the higher-order assembly samples. While the peak current maxima vs dwell time density scatter plots highlighted significant differences between samples, to extract more detail from the translocation information, we investigated an additional nanopore translocation discriminant. The observed translocation peaks are conductive, and therefore imply that an increased amount of charge is passing through the nanopore during the translocation event compared to the baseline while no DNA origami pass through. We define the Equivalent Charge Surplus (ECS) of each translocation event as the area of the conductive translocation peak (Figure 6A) (37).

The ion current signature discriminants used above (peak current maximum and translocation dwell time) are likely dependent on the shape and orientation of the nanostructure during translocation through the nanopore. In contrast, the overall charge is expected to remain conserved for the same higher-order assembly state of DNA origami, and can be expected to scale linearly with the size of the higher-order assemblies made from identical DNA origami (monomers). This is confirmed in Figure 6B, where the average ECS values of the DNA origami designs are plotted versus their surface area. The red line represents a linear fit to the data. Figure 6C depicts the ECS distributions for each higher-order DNA origami sample analysed with the nanopore platform. In contrast to the distributions of the peak current maxima (Figure 4C), even for the most complex sample (2×2 sample) four clearly discernible peaks can be seen in the distribution, which suggests that each peak corresponds to either the monomer building block, the fully assembled structure, or a particular higher-order assembly intermediate. To allocate the different distribution peaks to a particular structure, we used the information obtained from less complex assemblies in the analysis of the more complex assemblies. Clearly, the monomer sample is expected to contain predominantly monomers and at most a negligible amount of non-specifically assembled higher-order structures. This is reflected in the very prominent peak centred around an ECS of ∼150 pC (left-most panel of Figure 6C). The solid line represents a Gaussian fit to the distribution, which yielded an average ECS of 143 pC. The barely noticeable contribution to the distribution at around 300 pC may indicate a negligible presence of non-specific assemblies, but the contribution is difficult to quantify and is significantly smaller. Therefore, we allocate the ECS peak centred at 143 pC (labelled I) to the monomer. The ECS distribution of the dimer sample shows two peaks (second panel of Figure 6C). In this assembly sample we expect the monomer and of course the dimer to be present. Therefore, we fitted the distribution with a two-peak Gaussian while keeping the first peak fixed at 143 pC, the position obtained from the monomer sample (most-left panel of Figure 6C). The fits are represented by solid lines in the figure; the red line represents the fixed monomer peak (I), and the blue line the additional peak (II) which yielded an average ECS of 332 pC, and which we attribute to the presence of the dimer. To confirm that peak II indeed represents the dimer, we sliced the dimer sample events according to which Gaussian peak of the ECS distribution they belong, and generated the associated peak current maxima versus dwell time density scatter plots (Figure S11). As shown in the second row of Figure S11, the monomer component in Figure 5B is not present in the density scatter plot (panel C of Figure S11), supporting the viability of this ECS clustering to mark the different DNA nanostructure components present in the assembly samples. We then applied the same approach for the trimer and the 2×2 sample, resulting in an ECS distribution peak at 481 pC (III) which represents the trimer, and an ECS distribution peak at 636 pC (IV) which represents the 2×2 component. We note a slight discrepancy between the fixed monomer and dimer peaks in the ECS distributions compared to the observed distribution for both the trimer and 2×2. This is likely associated with small variations in pore size of the nanopipettes used, as each dataset was obtained with different nanopipettes. However, the deviations are small and do not impact on our ability to distinguish different populations. The equivalent data slicing as for the dimer sample was carried out and, like the dimer samples, the fully assembled DNA nanostructures yielded well-defined isolated clusters in the respective density scatter plots (Figure S11C). While all current traces were recordings of 3 min, we carried out the same analysis on a shorter trace (1 min) with correspondingly fewer translocation events, and we show in Figure S12 that the results remain consistent for shorter current traces (1min vs 3min recorded trace).

The areas of the Gaussian fits to the ECS distribution peaks allow us to associate individual translocation event to a distinct assembly state (monomer building block, fully assembled DNA nanostructure, or assembly intermediate) and thus to estimate the percentage of each higher-order DNA nanostructure present in the samples and compute an assembly yield for each DNA construct. Figure 6E (blue bars) shows the yield for forming the desired end-product for each higher-order assembly sample, *i*.*e*. the percentage of monomer (I), dimer (II), trimer (III) and 2×2 (IV) in the monomer, dimer, trimer and 2×2 sample, respectively. As expected from the histogram in Figure 6C, the yield for the monomer is 100%, while yields for the higher-order assemblies decrease with the increase in assembly size (all numerical yield values are supplied in Table S1).

Traditionally, agarose gel electrophoresis is often employed to estimate assembly yields. An electrophoresis gel containing lanes for each higher-order assembly is shown in Figure 6D, and the assembly yields were determined from the agarose gel by densitometry and taking into account the size of the structure. The results are shown as red bars in Figure 6E and the numerical values are shown in Table S1. Overall, the yields obtained with gel electrophoresis show the same trend as the ones obtained from nanopore measurements. However, there are some notable differences. The agarose gel indicated a larger percentage of monomers in the higher-order assemblies for the trimer and 2×2 sample compared to the nanopore measurements (Table S1). This could be a result of the long run time of gel electrophoresis (2 hrs) that may compromise the stability of the larger assemblies. Furthermore, the quantification via gel electrophoresis assumes that the fluorescent signal is proportional to the number of basepairs in the assembled structure, which may be an over-simplification. Another often employed means of assessing DNA origami structures is AFM. Our AFM micrographs (Figure S3) confirmed the heterogenous character of the higher-order DNA origami samples. AFM is extremely well suited to study small numbers of DNA origami at very high detail so a good understanding of their structures and fine detail of the folding can be obtained. However, their use for quantification of the assembly yield is questionable in this case. AFM scans depend heavily on the mica surface preparation which is required for imaging, and this is likely to lead to an under-representation of smaller DNA constructs owing to the higher sedimentation rate of the larger DNA origami structures onto the mica substrate. This was confirmed here, where we observed an under-representation of monomers in the higher-order assembly samples (trimer and 2×2 samples) compared to the nanopore measurements.

Compared to the standard analysis methods of AFM and agarose gel electrophoresis, the nanopore measurements offer several advantages. Our nanopore method allowed us to obtain a label-free analysis of the DNA origami samples within a few minutes with single molecule resolution at statistically relevant numbers, and no lengthy sample preparations or use of expensive reagents. Another advantage of the nanopore approach is its single molecule analysis which can potentially detect minute concentrations and reveal the presence of DNA constructs that have formed with very low yields. In Figure S8F we demonstrate nanopore detection of DNA origami nanostructures down to the 50 pM concentrations.

In addition to utilising nanopore measurements for the determination of assembly yields of DNA origami, the approach enables a range of other applications. For example, the ability to differentiate between assembly states enables the probing of the association/dissociation of higher-order DNA constructs and shed light on their stability in different assembly configurations. Furthermore, the analysis is real-time and non-destructive, *i*.*e*. the DNA origami nanostructures could be collected in the bath after translocation and reused. This opens up the possibility for using the nanopore approach in label-free separation or purification, where, depending on the translocation peak characteristics, the DNA nanostructure can be steered (*e*.*g*. electrophoretically) into a collection or waste tube.

## CONCLUSIONS

In conclusion, we explored nanopore translocation as a single molecule approach to probe the heterogeneous character of DNA origami assemblies. The large number of events which can be recorded (>1000 events) for each sample within minutes enables statistically relevant studies in a non-destructive and label-free way. We demonstrated the discrimination of various assembly states for higher-order DNA origami assemblies based on their equivalent charge surplus computed from the recorded ion current signals, which allowed the quantification of the assembly yields without any lengthy sample preparations, and importantly enables a range of other applications where rapid single-molecule detection is required. Our work complements related approaches of using nanopore translocations characteristics to differentiate between DNA nanostructures with different geometries (41-43), but in contrast to most published studies on nanopore analysis of DNA origami, here we expand the scope of the approach to large higher-order assemblies.

## Author Contributions

SC performed the nanopore sensing, AFM measurements, and analysed the data. GM performed nanopore sensing experiments and analysed the data. IS designed and formed the DNA nanostructures and performed AFM measurements. CW and PA supervised the research and supported the data analysis. All authors wrote the manuscript.

## Declaration of Interests

The authors have no interests to declare.

## Acknowledgements

We thank Dr. Alexander Kulak for the help provided with SEM imaging of the nanopipettes. S.C. and P.A. acknowledge funding from the European Union’s Horizon 2020 research and innovation 7 program under the Marie Skłodowska-Curie MSCA-ITN grant agreement no. 812398, through the single-entity nanoelectrochemistry, SENTINEL, project. I.S. was supported by funding from the European Research Council (ERC) under the project DYNAMIN, grant agreement no. 788968. G.M. acknowledges funding from a University of Leeds Scholarship. P.A and C.W. acknowledge funding form the Engineering and Physical Science Research Council UK (EPSRC) Healthcare Technologies for the grant EP/W004933/1, and C.W. acknowledges funding from the Medical Research Council (MRC) UK under the grant number MR/N029976/1. Schematics used in our figures were adapted from BioRender.com.

## Notes

### Competing Interest Statement

The authors have declared no competing interest.

## REFERENCES

1. Seeman, N.C. and Sleiman, H.F. DNA nanotechnology. Nature Reviews Materials. 2017, 3(1), p.17068.

2. Dey, S., Fan, C., Gothelf, K.V., Li, J., Lin, C., Liu, L., Liu, N., Nijenhuis, M.A.D., Saccà, B., Simmel, F.C., Yan, H. and Zhan, P. DNA origami. Nature Reviews Methods Primers. 2021, 1(1), p.13.

3. Chandrasekaran, A.R., Anderson, N., Kizer, M., Halvorsen, K. and Wang, X. Beyond the Fold: Emerging Biological Applications of DNA Origami. ChemBioChem. 2016, 17(12), pp.1081–1089.

4. Engelhardt, F.A.S., Praetorius, F., Wachauf, C.H., Bruggenthies, G., Kohler, F., Kick, B., Kadletz, K.L., Pham, P.N., Behler, K.L., Gerling, T. and Dietz, H. Custom-Size, Functional, and Durable DNA Origami with Design-Specific Scaffolds. ACS Nano. 2019, 13(5), pp.5015–5027.

5. Zhang, Y., Li, Q., Liu, X., Fan, C., Liu, H. and Wang, L. Prescribing DNA Origami Patterns via Scaffold Decoration. Small. 2020, 16(16), p.e2000793.

6. Tikhomirov, G., Petersen, P. and Qian, L. Fractal assembly of micrometre-scale DNA origami arrays with arbitrary patterns. Nature. 2017, 552(7683), pp.67–71.

7. Wang, X., Jun, H. and Bathe, M. Programming 2D Supramolecular Assemblies with Wireframe DNA Origami. Journal of the American Chemical Society. 2022, 144(10), pp.4403–4409.

8. Gu, H., Chao, J., Xiao, S.-J. and Seeman, N.C. Dynamic patterning programmed by DNA tiles captured on a DNA origami substrate. Nature Nanotechnology. 2009, 4(4), pp.245–248.

9. Huang, Y., Nguyen, M.-K., Natarajan, A.K., Nguyen, V.H. and Kuzyk, A. A DNA Origami-Based Chiral Plasmonic Sensing Device. ACS Applied Materials & Interfaces. 2018, 10(51), pp.44221–44225.

10. Arroyo-Currás, N., Sadeia, M., Ng, A.K., Fyodorova, Y., Williams, N., Afif, T., Huang, C.-M., Ogden, N., Andresen Eguiluz, R.C., Su, H.-J., Castro, C.E., Plaxco, K.W. and Lukeman, P.S. An electrochemical biosensor exploiting binding-induced changes in electron transfer of electrode-attached DNA origami to detect hundred nanometer-scale targets. Nanoscale. 2020, 12(26), pp.13907–13911.

11. Selnihhin, D., Sparvath, S.M., Preus, S., Birkedal, V. and Andersen, E.S. Multifluorophore DNA Origami Beacon as a Biosensing Platform. ACS Nano. 2018, 12(6), pp.5699–5708.

12. Raveendran, M., Leach, A.R., Hopes, T., Aspden, J.L. and Actis, P. Ribosome Fingerprinting with a Solid-State Nanopore. ACS Sensors. 2020, 5(11), pp.3533–3539.

13. Raveendran, M., Lee, A.J., Sharma, R., Wälti, C. and Actis, P. Rational design of DNA nanostructures for single molecule biosensing. Nature Communications. 2020, 11(1), p.4384.

14. Zhang, Q., Jiang, Q., Li, N., Dai, L., Liu, Q., Song, L., Wang, J., Li, Y., Tian, J., Ding, B. and Du, Y. DNA Origami as an In Vivo Drug Delivery Vehicle for Cancer Therapy. ACS Nano. 2014, 8(7), pp.6633–6643.

15. Ge, Z., Guo, L., Wu, G., Li, J., Sun, Y., Hou, Y., Shi, J., Song, S., Wang, L., Fan, C., Lu, H. and Li, Q. DNA Origami-Enabled Engineering of Ligand-Drug Conjugates for Targeted Drug Delivery. Small. 2020, 16(16), p.e1904857.

16. Lee, A.J., Endo, M., Hobbs, J.K. and Wälti, C. Direct Single-Molecule Observation of Mode and Geometry of RecA-Mediated Homology Search. ACS Nano. 2018, 12(1), pp.272–278.

17. Lee, A.J., Endo, M., Hobbs, J.K., Davies, A.G. and Walti, C. Micro-homology intermediates: RecA’s transient sampling revealed at the single molecule level. Nucleic Acids Res. 2021, 49(3), pp.1426–1435.

18. Platnich, C.M., Rizzuto, F.J., Cosa, G. and Sleiman, H.F. Single-molecule methods in structural DNA nanotechnology. Chemical Society Reviews. 2020, 49(13), pp.4220–4233.

19. Xue, L., Yamazaki, H., Ren, R., Wanunu, M., Ivanov, A.P. and Edel, J.B. Solid-state nanopore sensors. Nature Reviews Materials. 2020, 5(12), pp.931–951.

20. Alfaro, J.A., Bohländer, P., Dai, M., Filius, M., Howard, C.J., van Kooten, X.F., Ohayon, S., Pomorski, A., Schmid, S., Aksimentiev, A., Anslyn, E.V., Bedran, G., Cao, C., Chinappi, M., Coyaud, E., Dekker, C., Dittmar, G., Drachman, N., Eelkema, R., Goodlett, D., Hentz, S., Kalathiya, U., Kelleher, N.L., Kelly, R.T., Kelman, Z., Kim, S.H., Kuster, B., Rodriguez-Larrea, D., Lindsay, S., Maglia, G., Marcotte, E.M., Marino, J.P., Masselon, C., Mayer, M., Samaras, P., Sarthak, K., Sepiashvili, L., Stein, D., Wanunu, M., Wilhelm, M., Yin, P., Meller, A. and Joo, C. The emerging landscape of single-molecule protein sequencing technologies. Nature Methods. 2021, 18(6), pp.604–617.

21. Shi, W., Friedman, A.K. and Baker, L.A. Nanopore Sensing. Analytical Chemistry. 2017, 89(1), pp.157–188.

22. Houghtaling, J., Ying, C., Eggenberger, O.M., Fennouri, A., Nandivada, S., Acharjee, M., Li, J., Hall, A.R. and Mayer, M. Estimation of Shape, Volume, and Dipole Moment of Individual Proteins Freely Transiting a Synthetic Nanopore. ACS Nano. 2019, 13(5), pp.5231–5242.

23. Bacri, L., Oukhaled, A.G., Schiedt, B., Patriarche, G., Bourhis, E., Gierak, J., Pelta, J. and Auvray, L. Dynamics of Colloids in Single Solid-State Nanopores. The Journal of Physical Chemistry B. 2011, 115(12), pp.2890–2898.

24. Karmi, A., Dachlika, H., Sakala, G.P., Rotem, D., Reches, M. and Porath, D. Detection of Au Nanoparticles Using Peptide-Modified Si3N4 Nanopores. ACS Applied Nano Materials. 2021, 4(2), pp.1000–1008.

25. Arima, A., Harlisa, I.H., Yoshida, T., Tsutsui, M., Tanaka, M., Yokota, K., Tonomura, W., Yasuda, J., Taniguchi, M., Washio, T., Okochi, M. and Kawai, T. Identifying Single Viruses Using Biorecognition Solid-State Nanopores. Journal of the American Chemical Society. 2018, 140(48), pp.16834–16841.

26. Yu, R.-J., Lu, S.-M., Xu, S.-W., Li, Y.-J., Xu, Q., Ying, Y.-L. and Long, Y.-T. Single molecule sensing of amyloid-β aggregation by confined glass nanopores. Chemical Science. 2019, 10(46), pp.10728–10732.

27. Engst, C.R., Ablay, M., Divitini, G., Ducati, C., Liedl, T. and Keyser, U.F. DNA Origami Nanopores. Nano Letters. 2012, 12(1), pp.512–517.

28. Wang, V., Ermann, N. and Keyser, U.F. Current Enhancement in Solid-State Nanopores Depends on Three-Dimensional DNA Structure. Nano Letters. 2019, 19(8), pp.5661–5666.

29. Plesa, C., van Loo, N., Ketterer, P., Dietz, H. and Dekker, C. Velocity of DNA during Translocation through a Solid-State Nanopore. Nano Letters. 2015, 15(1), pp.732–737.

30. Plesa, C., Ananth, A.N., Linko, V., Gülcher, C., Katan, A.J., Dietz, H. and Dekker, C. Ionic Permeability and Mechanical Properties of DNA Origami Nanoplates on Solid-State Nanopores. ACS Nano. 2014, 8(1), pp.35–43.

31. Raveendran, M., Lee, A.J., Wälti, C. and Actis, P. Analysis of 2D DNA Origami with Nanopipettes. ChemElectroChem. 2018, 5(20), pp.3014–3020.

32. Chau, C.C., Radford, S.E., Hewitt, E.W. and Actis, P. Macromolecular Crowding Enhances the Detection of DNA and Proteins by a Solid-State Nanopore. Nano Letters. 2020, 20(7), pp.5553–5561.

33. Beamish, E., Tabard-Cossa, V. and Godin, M. Programmable DNA Nanoswitch Sensing with Solid-State Nanopores. ACS Sensors. 2019, 4(9), pp.2458–2464.

34. Chen, K., Zhu, J., Bošković, F. and Keyser, U.F. Nanopore-Based DNA Hard Drives for Rewritable and Secure Data Storage. Nano Letters. 2020, 20(5), pp.3754–3760.

35. Tikhomirov, G., Petersen, P. and Qian, L. Programmable disorder in random DNA tilings. Nature Nanotechnology. 2017, 12(3), pp.251–259.

36. Douglas, S.M., Marblestone, A.H., Teerapittayanon, S., Vazquez, A., Church, G.M. and Shih, W.M. Rapid prototyping of 3D DNA-origami shapes with caDNAno. Nucleic Acids Research. 2009, 37(15), pp.5001–5006.

37. Plesa, C. and Dekker, C. Data analysis methods for solid-state nanopores. Nanotechnology. 2015, 26, p.084003.

38. Wanunu, M., Sutin, J., McNally, B., Chow, A. and Meller, A. DNA Translocation Governed by Interactions with Solid-State Nanopores. Biophysical journal. 2008, 95, pp.4716–4725.

39. Li, J. and Talaga, D. The distribution of DNA translocation times in solid-state nanopores. Journal of physics. Condensed matter : an Institute of Physics journal. 2010, 22, p.454129.

40. Zhu, L., Zhang, Z. and Liu, Q. Deformation-Mediated Translocation of DNA Origami Nanoplates through a Narrow Solid-State Nanopore. Anal Chem. 2020, 92(19), pp.13238–13245.

41. Xie, Z.-P., Liu, S.-m. and Zhai, Y.-M. Study on the Self-assembly and Signal Amplification Ability of Nucleic Acid Nanostructure with the Nanopipette. Journal of Electroanalytical Chemistry. 2022, 914, p.116307.

42. Yang, J., Zhao, N., Liang, Y., Lu, Z. and Zhang, C. Structure-flexible DNA origami translocation through a solid-state nanopore. RSC Advances. 2021, 11(38), pp.23471–23476.

43. Alibakhshi, M.A., Halman, J.R., Wilson, J., Aksimentiev, A., Afonin, K.A. and Wanunu, M. Picomolar Fingerprinting of Nucleic Acid Nanoparticles Using Solid-State Nanopores. ACS Nano. 2017, 11(10), pp.9701–9710.

