## Supporting Information for "Nanopore Fingerprinting of Supramolecular DNA Nanostructures"

### **S1: Nanopore characterization**

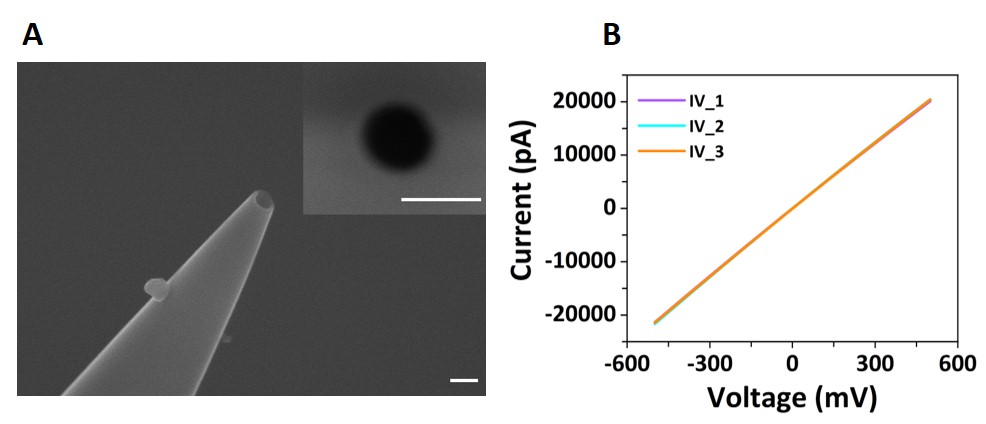

**Figure S1: (A)** SEM micrograph of a nanopipette used for translocation experiments, with a top view inset of the pore (200 nm scale bars). The diameter of the pore used in this study was approximately 160 nm. **(B)** Current-voltage curves (IV) of three nanopipettes recorded in 0.1M KCl.

### **S2: Agarose gel electrophoresis analysis of DNA nanostructures**

Quantification of the relative amounts of monomers, various assembly intermediates, and final products for each DNA origami sample was carried out by measuring the corresponding bands’ intensity with ImageJ and normalised by the size of the higher-order assemblies to account for the amount of double stranded DNA stained per macromolecule. As shown in Figure S2, the gel electrophoresis analysis confirmed the presence of multiple DNA nanostructures in the higher-order assembly samples, *i.e.* the dimer sample contains the monomer nanostructures as well. The trimer sample contains monomer and dimer nanostructures. The 2x2 samples contains monomer, dimer, and trimer nanostructures.

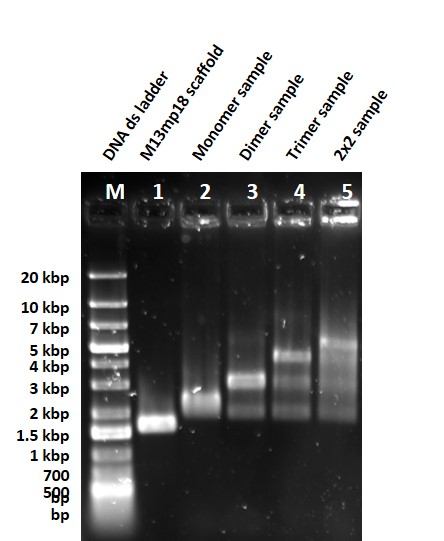

**Figure S2:** Agarose gel electrophoresis of DNA origami samples used in this study. Lane M: GeneRuler 1kb dsDNA ladder; Lane 1: M13mp18 circular ssDNA; Lane 2: monomer DNA origami sample; Lane 3: dimer DNA origami sample; Lane 4: trimer DNA origami sample; Lane 5: 2x2 DNA origami sample. Note, Figure 6D is a subset of this figure.

### **S3: AFM analysis of DNA origami samples**

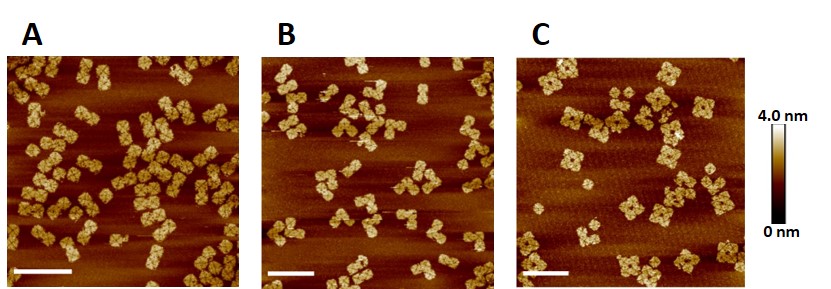

**Figure S3:** AFM micrographs of the higher-order assembly DNA origami samples: dimer **(A)**, trimer **(B)**, and 2x2 **(C)** using a 2 µm x 2 µm scan size (400 nm scale bars).

### **S4: Nanopore detection enhancement in PEG electrolyte bath**

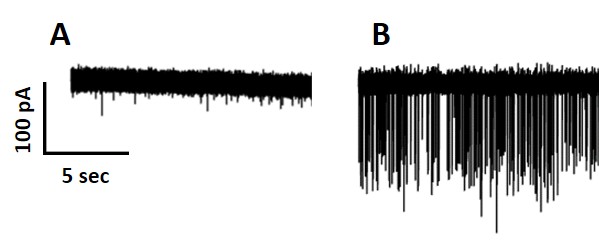

**Figure S4:** Ion current traces for the monomer DNA origami sample recorded in 0.1M KCl electrolyte bath **(A)** and 50% (w/v) PEG 35k 0.1M KCl bath **(B)**. Current traces were recorded at -300mV using the same nanopipette filled with monomer DNA nanostructures. The current and time scales are identical for both plots.

### **S5: Nanopore measurement stability**

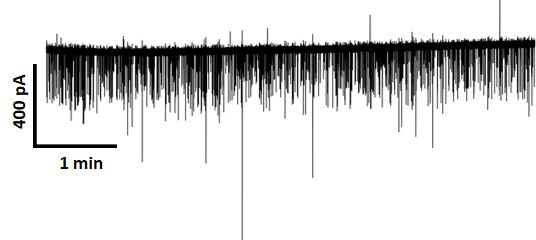

**Figure S5:** Ion Current trace of the 2x2 DNA origami sample recorded for 6 minutes under an applied voltage of -300mV.

### **S6: Translocation events detection from nanopore current traces**

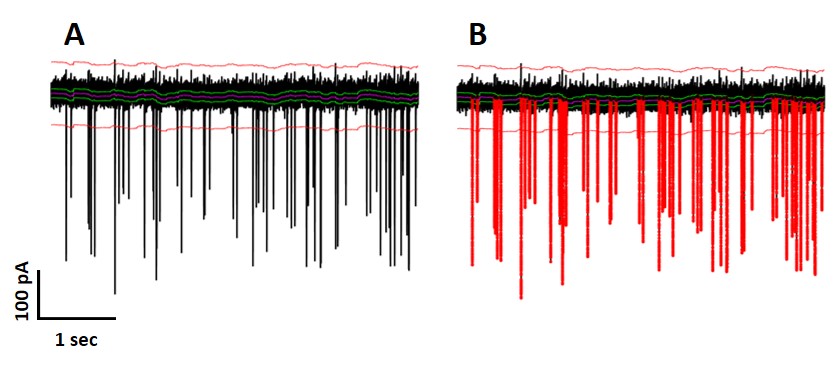

**Figure S6: (A)** Representative ion current trace with the 7σ threshold indicated by the red line. **(B)** Same trace as in A with the perturbations in the ion current identified as DNA origami translocation events by the Transanalyser Matlab script (1) according to the 7σ threshold highlighted in red.

### **S7: Translocation controls DNA nanostructures**

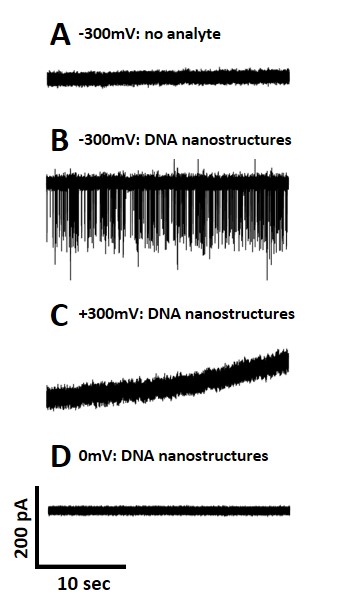

**Figure S7:** Control ion currents traces. **(A)** translocation at -300 mV when no analyte is added inside the nanopipette; **(B)** translocation at -300 mV when monomer DNA origami sample is added inside the nanopipette; **(C)** translocation at +300 mV when monomer DNA origami sample is added inside the nanopipette; **(D)** translocation at 0 mV when monomer DNA origami sample is added inside the nanopipette. The current and timescales are the same for all graphs.

### **S8: Translocation event characteristics DNA nanostructures**

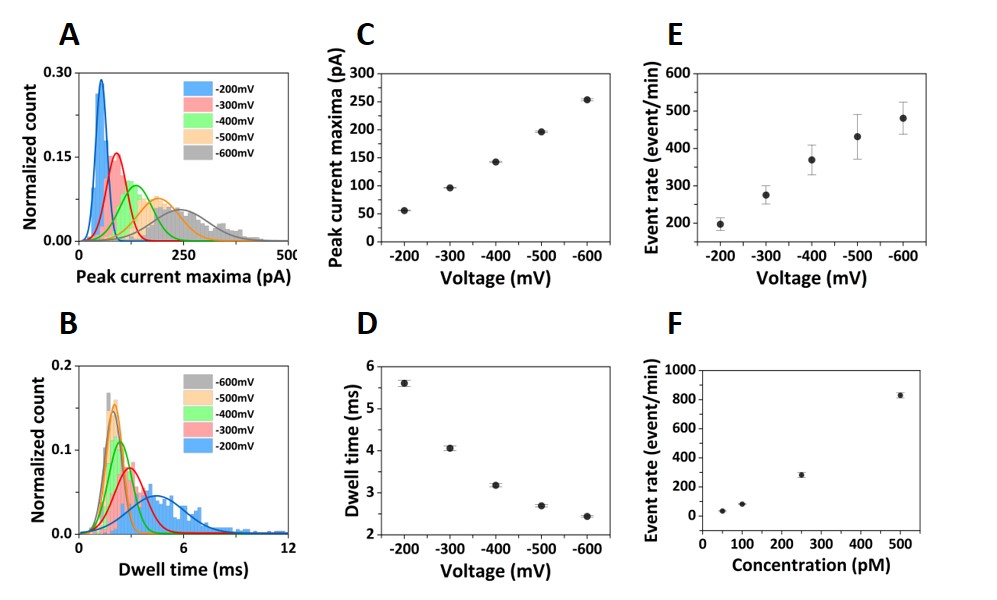

**Figure S8:** Histograms of the current peak maxima **(A)** and the dwell time **(B)** distributions for increasing absolute values of applied voltages *V* = -200 mV, -300 mV, -400 mV, -500 mV, and -600 mV. The solid lines represent Gaussian fits to the distributions. Average peak current maxima **(C)** and dwell time **(D)** at *V* = -200 mV, -300 mV, -400 mV, -500 mV, and -600 mV. Error bars show the standard deviation from three independent recordings. Event rate of the DNA nanostructures translocation as a function of *V* **(E)** and sample concentration (50 pM, 100 pM, 250 pM, and 500 pM) **(F)**. Error bars show the standard deviation from three independent recordings. Monomer DNA origami sample (250 pM for panel A-E) was used for these translocation experiments, and each translocation condition was repeated independently three times.

### **S9: Translocation events comparison for different DNA origami samples**

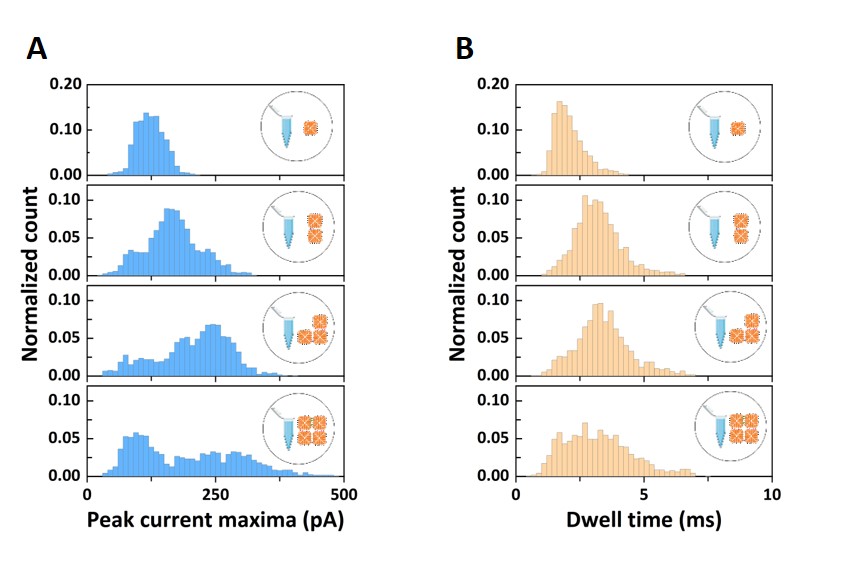

**Figure S9:** Histograms for peak current maxima **(A)** and dwell time **(B)** distributions for each DNA origami sample investigated. From top to bottom: monomer, dimer, trimer, and 2x2 samples. Translocation events were recorded at -300 mV.

### **S10: Monomer spiking**

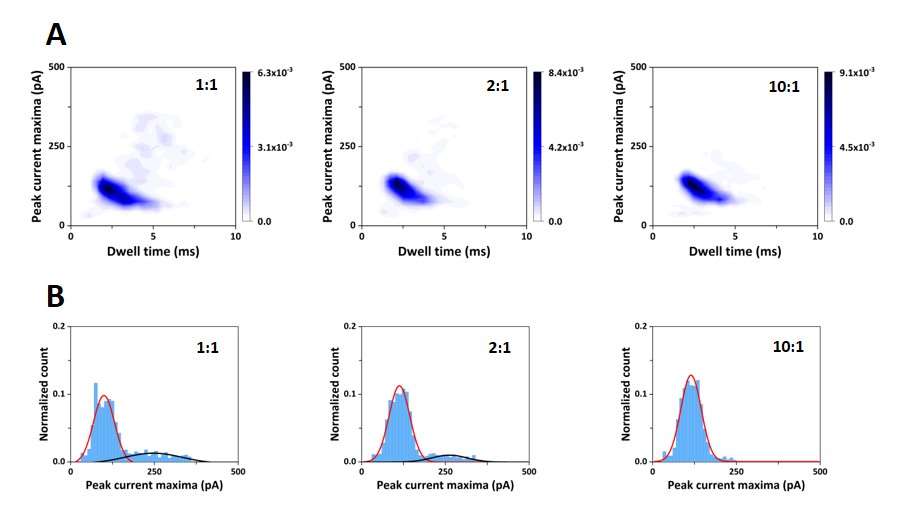

**Figure S10: (A)** Density scatter plots of the peak current maxima as a function of dwell time for 2x2 sample spiked with monomer sample. **(B)** Histograms of the current peak maxima distributions for the 2x2 sample spiked with monomer sample with multi-peak Gaussian fits represented by the solid lines. The notation 1:1, 2:1, and 10:1 for the left-to-right graphs in each panel indicates the molar ratio of monomer sample spiked into the 2x2 sample. Solid lines are Gaussian fits to the distributions.

### **S11: DNA nanostructures cluster isolation based on ECS**

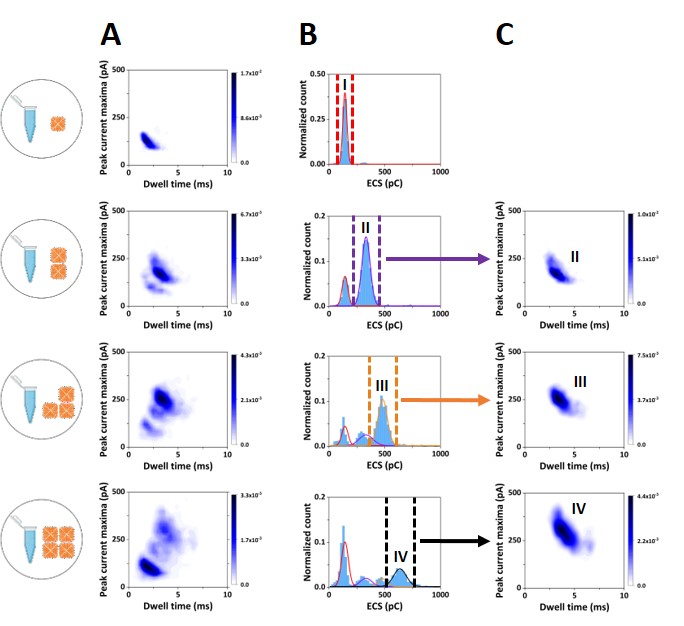

**Figure S11: (A)** Density scatter plots of the current peak maxima as a function of dwell time for (top to bottom): monomer sample, dimer sample, trimer sample, and 2x2 sample. **(B)** ECS histograms for the DNA origami samples presented in panel A. Marked Gaussian fitted peaks correspond to the DNA nanostructures of interest in each sample. **(C)** The corresponding sliced cluster depicted in a density scatter plot using only the translocation events sliced according to the marked Gaussian fitted peaks shown in Panel B. We note that the intensity scales are normalised to the number of translocation events shown, and therefore are different compared to panel A.

### **S12: Ratio of DNA nanostructures at different number of translocation events**

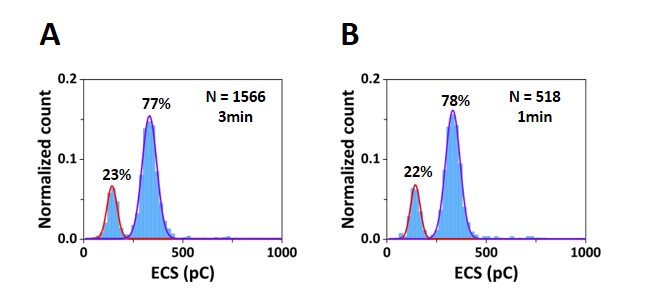

**Figure 12:** ECS histograms for dimer sample based on a 3-minute translocation recording **(A)** and 1-minute translocation recording **(B)**. Solid lines are Gaussian fits to the distributions.

### **S13: Ratio of DNA nanostructures nanopore *vs* gel electrophoresis**

**Table S1:** Ratio of DNA nanostructures determined for each DNA origami sample analyzed. The notation I-IV refers to the DNA origami nanostructures present in the sample analyzed: I - monomer nanostructure, II - dimer nanostructure, III - trimer nanostructure, and IV - 2x2 nanostructures. The dash indicates that the component was not detected and no ratio was computed.

| **DNA origami sample** | **Ratio DNA nanostructure nanopore analysis (%)** | | | | **Ratio DNA nanostructure agarose gel analysis (%)** | | | |
| --- | --- | --- | --- | --- | --- | --- | --- | --- |
|  | **I** | **II** | **III** | **IV** | **I** | **II** | **III** | **IV** |
| **Monomer Sample** | **100** | **-** | **-** | **-** | **100** | **-** | **-** | **-** |
| **Dimer Sample** | **23** | **77** | **-** | **-** | **12** | **88** | **-** | **-** |
| **Trimer Sample** | **17** | **21** | **62** | **-** | **48** | **19** | **33** | **-** |
| **2x2 Sample** | **44** | **14** | **10** | **33** | **66** | **13** | **3** | **18** |

### **S14: DNA nanostructures design**

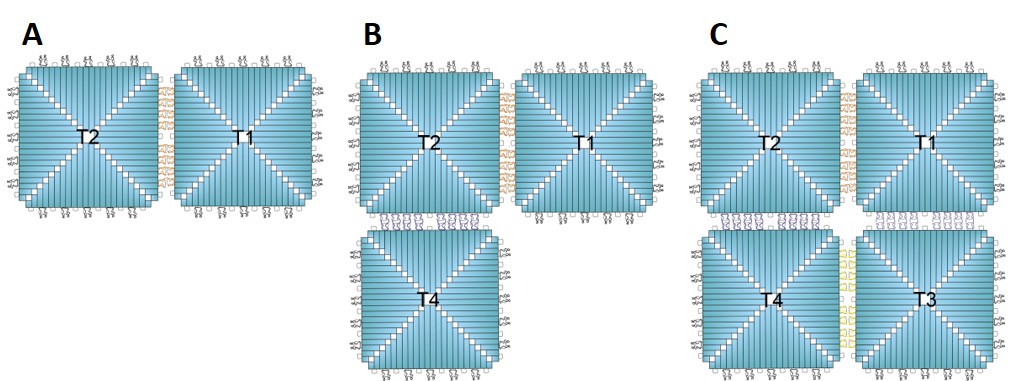

**Figure S14:** Designs of the higher-order assembly DNA nanostructures indicating the sets of monomer building-blocks used for their assembly: **(A)** - dimer, **(B)** - trimer, **(C)** - 2x2 nanostructure. The sequences for each set of staples are given in the excel file in SI. The notation T1-T4 refers to edge modifications on the monomer nanostructure in order to assemble the higher order DNA nanostructures.

### **References**

1.	Plesa, C. and Dekker, C. Data analysis methods for solid-state nanopores. *Nanotechnology.* 2015, **26**, p.084003.
